# Many weak and few strong links: The importance of interaction strength distributions for stabilising patterns in competition networks

**DOI:** 10.1101/2024.01.25.577181

**Authors:** Franziska Koch, Anje-Margriet Neutel, David K.A. Barnes, Korinna T. Allhoff

## Abstract

Ecological networks tend to contain many weak and only few strong interactions. Furthermore, interaction strengths are often arranged or patterned in ways that enhance stability. However, little attention has been given to the relation between the “many weak and few strong links” distribution and the stabilising effect of patterning. Here, we focus on the stabilising effect of hierarchy in bryozoan competition networks, and demonstrate that it critically depends on a skewed distribution of interaction strengths. To this end, we first show that, in line with many other ecological networks, the empirically derived interaction strengths in competition networks were characterised by a high level of skewness, with many weak and few strong links. Then, we analysed the relationship between the interaction strength distributions, hierarchy and stability by comparing theoretical competition matrices with different distributions of interaction strengths. We found that the full stabilising effect of hierarchy only appeared when we used skewed interaction strengths produced by a gamma distribution, but not in matrices built with uniform or half-normal distributions. This has important methodological implications, since theoretical studies often assume normal or uniform distributions to study ecological stability, and therefore might overlook stabilising mechanisms. We conclude that since skewed interaction strengths are a common feature of ecological networks, they can be expected to play an important role in the relation between structure and stability in living systems.

## 2 Introduction

In diverse communities, direct and indirect interactions between species form complex ecological networks. Their complexity makes it challenging to predict how assemblages, communities and whole ecosystems will react to environmental changes and disturbances (Montoya et al., 2006; Woodward et al., 2010; Strona and Lafferty, 2016; Barnes et al., 2021). Understanding how the strengths of interactions in ecological networks relate to system stability has therefore been a long-standing focus in the field of community and ecosystem ecology (Yodzis, 1981; Ruiter et al., 1995; McCann et al., 1998; Emmerson and Yearsley, 2004; Jacquet et al., 2016; Landi et al., 2018).

There is ample evidence of a characteristic distribution of many weak and few strong links in empirical studies on ecological networks, in particular in food webs (Paine, 1992; Neutel, Heesterbeek, and De Ruiter, 2002; Berlow, 1999; O’Gorman et al., 2010; Jacquet et al., 2016) but also in mutualistic systems (Jordano, 1987; Bascompte, Jordano, et al., 2006). Theoretical studies (McCann et al., 1998; Emmerson and Yearsley, 2004; James et al., 2015; Van Altena et al., 2016; Gellner and McCann, 2016; Jacquet et al., 2016) as well as some experimental evidence (O’Gorman and Emmerson, 2009) suggest that these weak links have a stabilising effect. Weak links contribute to stability by lowering the overall mean interaction strength (May, 1972), as well as through a dampening effect on oscillations, which has been well studied in small network modules (McCann et al., 1998; Emmerson and Yearsley, 2004). However, it has also been demonstrated that the particular arrangement (patterning) of strong and weak links has a large stabilising effect (Yodzis and Innes, 1992; Ruiter et al., 1995; James et al., 2015), which is often explained with respect to omnivorous loops (Neutel, Heesterbeek, and De Ruiter, 2002; Emmerson and Yearsley, 2004; Bascompte, Melián, et al., 2005; Wootton and Stouffer, 2016).

Thus, it is known that both the distribution of many weak and few strong links, and the patterning of link strengths are important for stability. This is in apparent contradiction to analytical results derived from random matrix theory. It has been shown that the exact shape of the distribution of interaction strengths does not influence the stability of large unstructured networks (Tao et al., 2010). Therefore, these studies typically use standard uniform or normal distributions (Allesina and Tang, 2012; Allesina and Tang, 2015), instead of more realistic, skewed distributions. For structured networks, however, it remains unclear how the distribution of interaction strengths influences the stabilising effect of patterning.

Here, we explore the relationship between stability, the interaction strength distribution and patterning in realistic, multi-species competitive communities. While it is obvious that some level of variation in strengths is needed to enable patterning, we do not know whether greater variation in link strengths will lead to stronger stabilising effects or how exactly the many weak few strong links distribution influences stability in patterned systems. We address these open questions using recent findings on the stabilising effect of competitive hierarchy in assemblages of encrusting, marine bryozoan colonies (Koch et al. (2023), see Box 1 for details). In networks with a competitive hierarchy, all species can be clearly ranked from strongest to weakest competitor. This strict ranking has been found to cause stabilising, asymmetric patterns both within pairs of competitors, and at the whole assemblage level. Koch et al. (2023) explain the stabilising effect of hierarchy based on the strength of feedback loops, closed chains of interactions that either amplify or dampen disturbances (see Fig. 1 in Box 1 for details).

**Figure 1:**
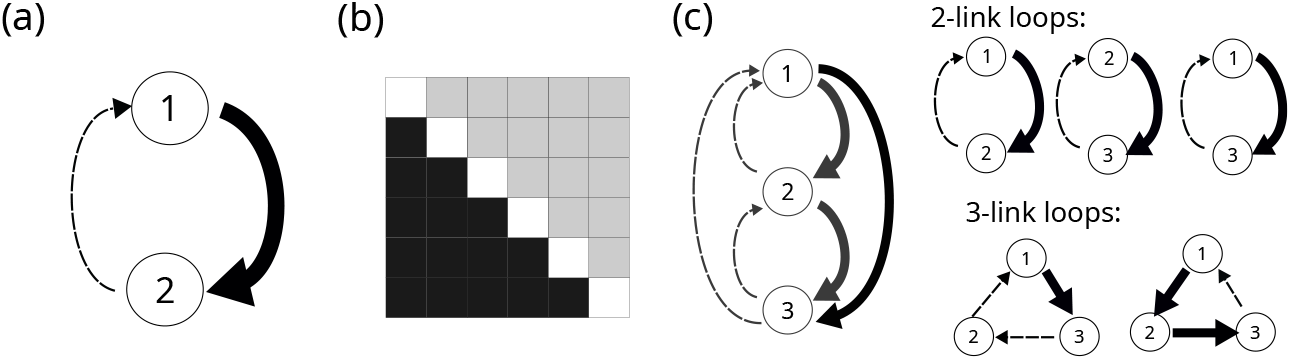
Asymmetric patterns in interaction strengths caused by hierarchical competition. (a) Pairwise asymmetry means that the loop formed by each pair of competing species consists of strong link, coupled to a much weaker link (b) Community asymmetry means that when we order the community matrix according to the hierarchical ranking, from strongest to weakest competitor, all strong links appear below the diagonal while all weak links appear above the diagonal. (c) Hierarchy reduces the strength of feedback loops, as each loop contains at least one weak link.

In a first step, we show that the “many weak, few strong” distribution that has been found in other network types also appears in competitive systems. Then, we ask if and to what extent the stabilising effect of hierarchy depends on this underlying distribution of link strengths. To address this question, we generated theoretical matrices in which we varied the proportion of strong and weak links by using random interaction strengths drawn from distributions with different shapes. For each type of distribution, we quantified the stabilising effect by comparing the stability of asymmetric matrices to unstructured ones, where links were randomly arranged (see Fig. 2 for a visual overview of our approach). We find that the stabilising effect of hierarchy critically depends on the skewness of the interaction strengths. This suggests that the “many weak, few strong” link distribution plays a critical role in stabilising ecosystems.

**Figure 2:**
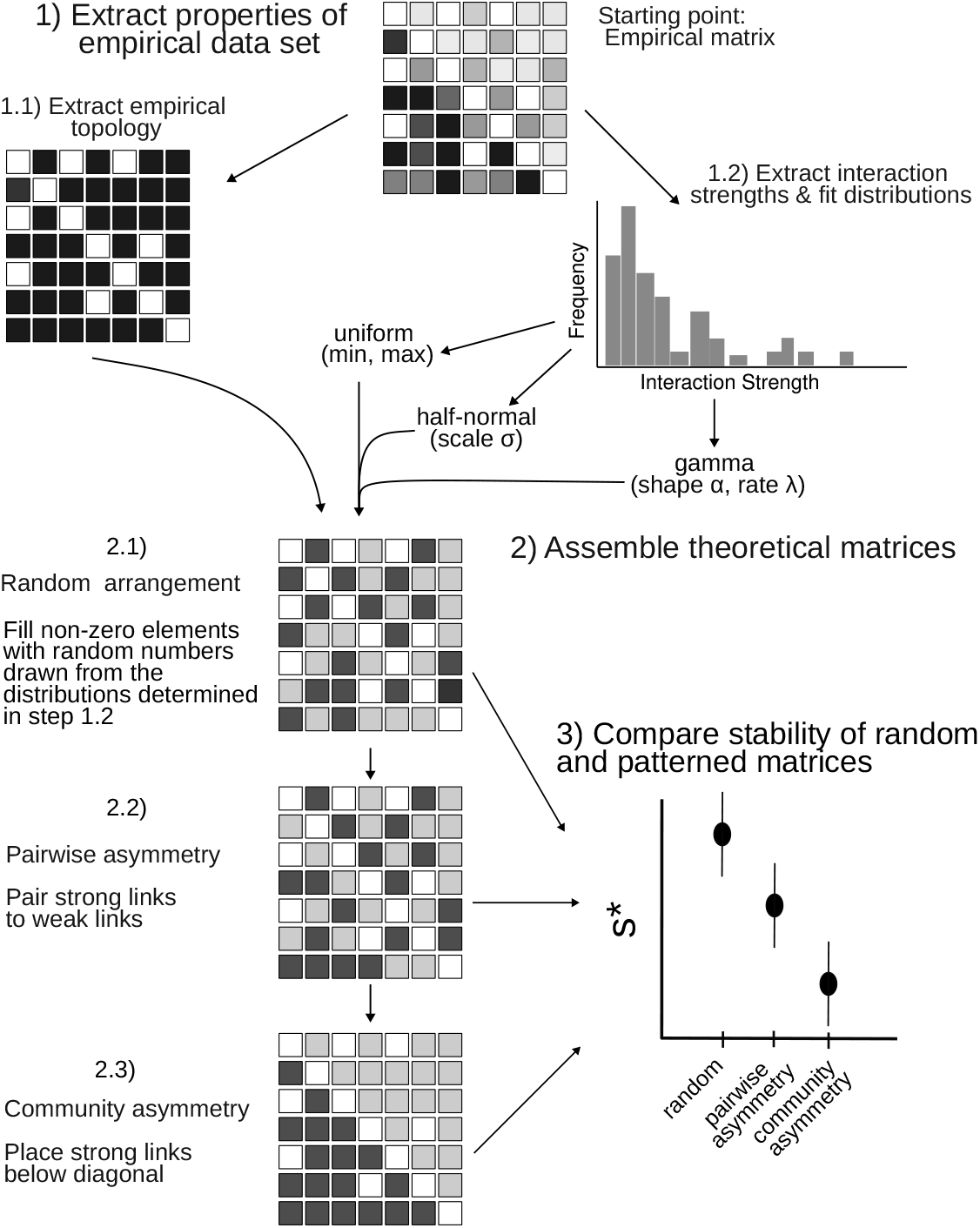
Assessing the relation between interaction strength distribution and stabilising effects of hierarchy using theoretical matrices. We started with a community matrix derived from empirical data, which contained normalised interspecific interactions of varying strengths (indicated by different shades of grey). Due to the normalisation, diagonal matrix elements, which represent intraspecific interactions, are equal to −1 (shown here in white for simplicity). As a first step (1.1), we extracted the empirical topology, the location of missing links (white off-diagonal cells) and we fitted uniform, half-normal and gamma distributions to the empirical interaction strengths (1.2). In a second step, the theoretical matrices were assembled using random interaction strengths generated with the fitted distributions. We then imposed three types of patterns on the theoretical matrices by arranging the theoretical interaction strengths in random (2.1), pairwise asymmetric (2.2) and community asymmetric (2.3) patterns. We preserved the locations of non-zero links extracted from the empirical matrix in this step. In a third step, we compared the stability of random, pairwise asymmetric and community asymmetric matrices. We measured stability as the critical amount of self-regulation needed for stability, *s*^*^. Lower values of *s*^*^ indicate more stable systems

### Box 1

**Explaining the stabilising effect of asymmetry in bryozoan assemblages (Koch et al., 2023)**

Koch et al. (2023) analysed the stability of 30 bryozoan assemblages by deriving inter- and intraspecific interaction strengths from observations of overgrowth competition between bryozoan colonies. They found that these interference competition networks had a hierarchical structure so that all species could be ranked from weakest to strongest competitor. This hierarchy results in asymmetric patterns, both within pairwise interactions, where a strong link is coupled with a weak link (Fig. 1 a), and at the community level, where strong links are concentrated on one side of the matrix diagonal (Fig. 1 b). While the observed competition networks are found to be unstable, their instability is reduced compared to randomised systems. This stabilising effect of asymmetric patterns of interaction strengths can be explained based on the concept of feedback loops, that is closed chains of interactions that connect one network element back to itself (Levins, 1974). They determine how perturbations propagate through the nodes and links of a network, through both direct and indirect effects (Zelnik et al., 2024). A loop is quantified as the product of all interaction strengths within the loop. Positive feedback loops (those with a positive product, for example 2-link loops in competitive systems) amplify disturbances that are introduced into the system, while negative feedback loops (those with a negative product, for example 3 link loops in competitive systems) counteract them (Levins, 1974).

Using an approach that was developed for trophic networks (Neutel, Heesterbeek, and De Ruiter, 2002; Neutel, Heesterbeek, Koppel, et al., 2007; Neutel and Thorne, 2014) and extending it for competition networks, Koch et al. (2023) show that the asymmetric patterns of interaction strengths reduce network instability by keeping short and long feedback loops weak. Pairwise asymmetry reduces instability of empirical competition networks by weakening the effect of positive 2-link feedback loops. As the effect of a loop depends on the product of link strengths, pairwise asymmetry means that a strong link is multiplied with a weak link (Fig. 1 c), so that the overall product remains low. The stabilising effect of community asymmetry, on the other hand, is related to longer, negative loops, which can cause unstable oscillations (Levins, 1974). Community asymmetry avoids the formation of such long negative loops, as all longer loops also contain at least a single weak link, which keeps the overall effect of the loops low (Fig. 1).

## 3 Methods

### 3.1 Empirical interaction strength distributions

We obtained empirical distributions of interaction strengths from Koch et al. (2023). For background on the methodology used by Koch et al. (2023), we describe the procedure of deriving empirical interaction strengths here. Following May (1972), interaction strengths are defined as the elements of the Jacobian or “community matrix”, which contains the partial derivatives of an underlying system of differential equations, evaluated at equilibrium. The elements describe the per-capita effect [dimension 1*/t*] of a change in the biomass of species *j* on the biomass of species *i*. Koch et al. (2023)’s calculation assumes Lotka-Volterra equations at equilibrium, with the observed abundances corresponding to equilibrium densities. First, energy flow rates are estimated from observed outcomes of spatial contests between bryozoan colonies. Then, these energy flow rates are combined with the observed abundances to translate them to interaction strengths (see Supplementary Note 1 for a more detailed description). Finally, the interaction strengths are normalised by dividing all off-diagonal terms by their corresponding diagonal term (following Neutel and Thorne, 2014). This scaling procedure preserves the essential stability characteristics (see Thorne et al. (2021) for a detailed explanation) and makes the matrices dimensionless, which simplifies the comparison of stability (see section 3.2.3 *“Calculation of network stability via the critical amount of self-regulation”*).

As a starting point of our analysis, we describe the shapes of the empirically based normalised distributions of interaction strengths observed by Koch et al. (2023) by calculating mean, variance, minimum, maximum, as well as skewness and kurtosis (using the R-package *“moments”*) of the non-zero interspecific interaction strengths in each of the 30 empirical matrices.

### 3.2 Using theoretical matrices to quantify the stabilising effect of asymmetry under different distributions of interaction strengths

We used theoretical competition matrices with random interaction strengths to explore the stabilising effect of pairwise and community asymmetry under varying distributions of interaction strengths (see Fig 2 for a visual overview of our approach). Our goal was to create theoretical matrices that closely resembled the empirical systems, but differed in their proportion of weak and strong links. For each empirical data set, we extracted the empirical topology, that is the network size, connectance and the location of non-zero links (Fig. 2 step 1.1). Then, we fitted distributions with different shapes to the empirical interaction strengths (Fig. 2 step 1.2). These were then used to generate artificial interaction strengths (Fig. 2 step 2, see section *“Theoretical interaction strength distributions”* for details). We arranged the artificial interaction strengths within the theoretical matrix to form random, pairwise asymmetric and community asymmetric pattern (steps 2.1-2.3 in Fig. 2, see section *“Patterning of interaction strengths in theoretical matrices”* for details). We quantified the stabilising effect of asymmetric pattern by comparing the stability of a set of random matrices to a set of patterned matrices (Fig. 2 step 3).

As a frame of reference that allowed us to compare the behaviour of the theoretical matrices to the empirical systems, our analysis also included an additional set of matrices using the observed, empirical interaction strength distributions but arranged in the same way as the theoretical interaction strengths.

#### 3.2.1 Theoretical interaction strength distributions

The off-diagonal matrix elements, representing interspecific interaction strengths, were randomly drawn from probability distributions. Our goal was to obtain distributions with comparable ranges, but with different proportions of weak and strong values across this range. To achieve this, we first fitted half-normal, uniform and gamma distributions to each empirical data set using the *“fitdistrplus”* R-package (Delignette-Muller and Dutang, 2015). Then, we used the parameters of these fitted distributions (uniform: min and max; half-normal: scale *σ*, gamma: shape *α* and rate *λ*) to generate random interaction strengths. As the gamma and half-normal distribution are defined for positive values, we fitted to the absolute values of the interaction strengths. As our goal was to preserve the range of values, we could not preserve the mean interaction strengths in the half-normal and uniform distributions. For uniform distributions, the means were on average 3.6 times higher than the empirical mean. The means of the half-normal distributions were on average 1.5 times higher than the empirical means, while there was no difference between the means of the gamma distributions and the means of the empirical distributions.

The diagonal elements of each matrix, which represent intraspecific interaction strengths, were set to -1. We did this to make them comparable to the normalised, empirical matrices. It is also consistent with previous theoretical work using random matrices (May, 1972; Allesina and Tang, 2012), which thus implicitly use the assumption that the matrix elements represent normalised interaction strengths.

#### 3.2.2 Patterning of interaction strengths in theoretical matrices

We then imposed three types of patterns on the theoretical matrices: random, pairwise asymmetric and community asymmetric (following Koch et al., 2023). To preserve empirical topology, we placed random interaction strengths only in locations that also had non-zero link strengths in the original empirical matrix. We also created matrices with random topologies, to test the generality of our results (Supplementary Fig. 1).

In the random, unstructured matrices used for the main analysis (Fig. 2, step 2.1), each link strength was independently placed in a random position within the matrix. These matrices served as a null model, that allowed us to quantify the stabilising effect of asymmetry. To impose pairwise asymmetry (Fig. 2, step 2.2), we made sure that within each pair, a strong link was paired with a weak link. This was achieved by sorting the set of random link strengths from weakest to strongest. The weakest link was then paired with the strongest link, the second strongest with the second weakest, etc. The location within the matrix was assigned randomly. To impose community asymmetry (Fig. 2 step 2.3), we additionally controlled the positions above or below the matrix diagonal, so that all strong links were located below the matrix diagonal.

#### 3.2.3 Calculation of network stability via the critical amount of self-regulation

The theoretical competition matrices were assumed to represent Jacobian matrices of some underlying (potentially non-linear) system of differential equations, which has been linearised around an equilibrium point (May, 1972). System stability here is the local asymptotic stability of this equilibrium point, which describes whether a system has the ability to return to its steady state after an infinitesimally small disturbance. Traditionally, the relative stability is measured by comparing *Re*(*λ*_*d*_) of the Jacobian matrix, which in case of stability (*Re*(*λ*_*d*_) *<* 0) is called the resilience of a system (the speed of return to steady state). However, this relative stability can be generalized to encompass also the region where systems are unstable (*Re*(*λ*_*d*_) *>* 0). This broader quantification of relative stability is used here (see Koch et al., 2023).

In general, any unstable system can be made stable by artificially increasing the diagonal matrix elements (the intraspecific interaction strengths), while any stable system can be made unstable by decreasing the diagonal matrix elements. The critical amount of self-regulation uses this mechanism to measure stability. It is defined as the factor by which the observed intraspecific interaction strengths have to be multiplied to bring the matrix to the threshold between stability and instability (Neutel, Heesterbeek, and De Ruiter, 2002). In the case of an unstable system, *s*^*^ is larger than 1 and its value describes how much more self-regulation needs to be added to make the system stable. In the case of a stable system, *s*^*^ *<* 1, it describes the “buffering capacity” of a system.

As all matrices used in this study have uniform diagonals (all *a*_11_ = −1, see section 3.2.1), the critical amount of self-regulation (*s*^*^) is equivalent to *Re*(*λ*_*d*_) of the same matrix but with the diagonal set to 0 (for details see Supplementary Material of Neutel and Thorne, 2014). Hence, we could have used the eigenvalue as a stability metric here. However, *s*^*^ offers a more general way to quantify relative stability that also works for matrices with varying diagonal terms (Neutel, Heesterbeek, and De Ruiter, 2002). This is usually the case when deriving interaction strengths from empirical data, where intraspecific competition can differ a lot between species in a given community or assemblage. In order to emphasise our focus on stabilising patterns in realistic systems and to enable comparison to the results of Koch et al. (2023), we therefore use the critical amount of self-regulation instead of *Re*(*λ*_*d*_).

## 4 Results

### 4.1 Empirical interaction strengths can be closely approximated by a gamma distribution

We analysed the shapes of the distributions of the interaction strengths from 30 empirical competition networks published in Koch et al. (2023). Interaction strengths were the normalised elements of the Jacobian matrix (see Methods and Supplementary Note 1). For each of the 30 data sets, we calculated not only the mean and variance but also the skewness (*Ŝ*) and kurtosis 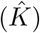 (Supplementary Table 1). Skewness describes a distribution’s asymmetry, while kurtosis describes how peaked a distribution is (Cristelli et al., 2012; Gross et al., 2021). We found that overall, the empirical interaction strengths showed distributions that were very asymmetric (mean *Ŝ* = −2.4) and peaked (mean 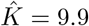), with a large variability in their skewness and kurtosis values (Fig. 3 a).

**Figure 3:**
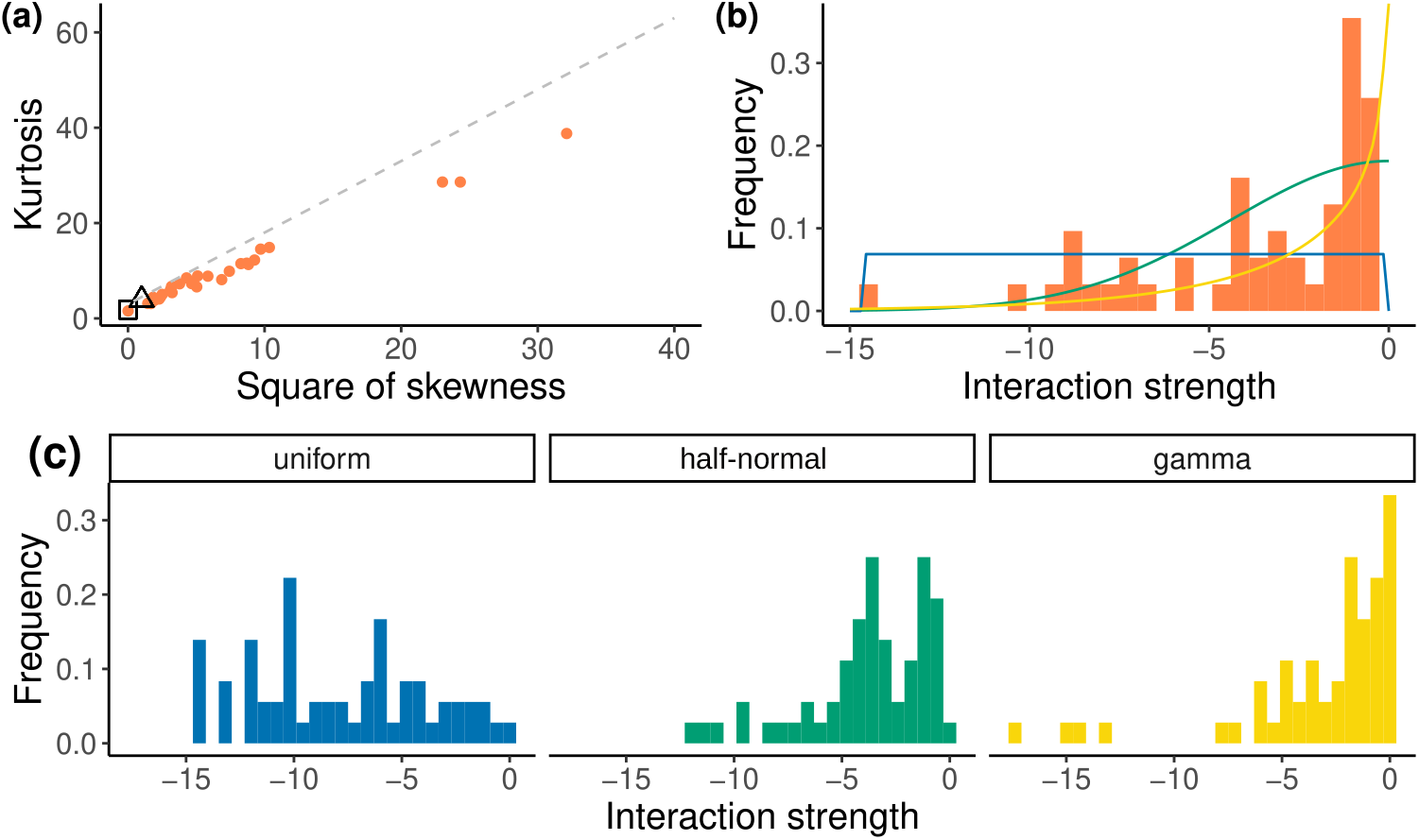
Skewed empirical interaction strengths are best represented by a gamma distribution. (a) Skewness-kurtosis relationship of the empirical distributions of interaction strengths on a Cullen and Frey graph, which can be used to differentiate between different types of distributions (Delignette-Muller and Dutang, 2015). Each orange dot represents one empirical data set. Normal and uniform distributions both only have one possible skewness *Ŝ* and kurtosis 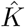 value (half-normal: *Ŝ* = 0.9, 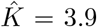, shown as a black triangle; uniform: *Ŝ* = 0, 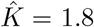 shown as a black square). The gamma distribution allows for varying *Ŝ* and 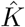 values and is represented as a dashed line, that represents how *Ŝ* and 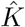 depend on the shape parameter. b) Histograms showing the empirical distribution of interaction strength for one example matrix (Rothera 3) as well as a uniform (blue), half-normal (green) and gamma (yellow) distribution fitted to these empirical values. (c) Example sets of random interaction strengths drawn from these fitted theoretical distributions.

The shapes of these empirical distributions thus differ clearly from half-normal and uniform distributions, which are more symmetric (*Ŝ* = 0.96 and *Ŝ* = 0, respectively) and have low kurtosis (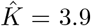 and 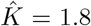). A gamma distribution (shown as the dotted line in Fig. 3 a) allows for varying skewness and kurtosis values and is able to capture the shape of empirical distributions much better. Fitting a gamma distribution to empirical interaction strengths allowed us to generate random interaction strengths with a more realistic pattern of many weak and few strong values (shown in Fig.3 b for one example data set). In contrast to that, a half-normal distribution produced too many intermediate values, while strong values were too rare. A uniform distribution, where all values are equally likely, generated too many strong values and too few weak ones (Fig. 3 c).

### 4.2 The stabilising effect of community asymmetry depends on the distribution of interaction strengths

We tested whether and how the distribution of link strengths affected the stabilising effect of asymmetries by comparing asymmetric to randomised theoretical competition matrices with uniform, half-normal and gamma shaped distributions of interaction strengths. For each type of distribution, we compare the stability (measured as critical self-regulation *s*^*^) of randomly patterned matrices to asymmetric ones, where strong links were paired to weak links as well as to community asymmetric ones, where strong links were also paired to weak links and all strong links additionally appear on one side of the diagonal (see Methods and Fig. 2).

Almost all theoretical matrices had critical amounts of self-regulation (*s*^*^) larger than 1 (Fig 4 a), indicating that the networks were unstable, and that additional self-regulation would be required to reach stability (where *s*^*^ = 1, or equivalently, *Re*(*λ*_*d*_) = 0). This was no surprise, as the random interaction strengths were fitted to link strengths from empirical community matrices that were also unstable (Koch et al., 2023). Throughout this analysis, we were thus comparing different levels of instability. Similarly, also the differences in instability between randomly patterned matrices of different distributions were expected (Fig. 4 a, light boxes). They arise from differences in mean interaction strengths (May, 1972): As the uniform distributions contained a higher proportion of strong values, these matrices were more unstable.

**Figure 4:**
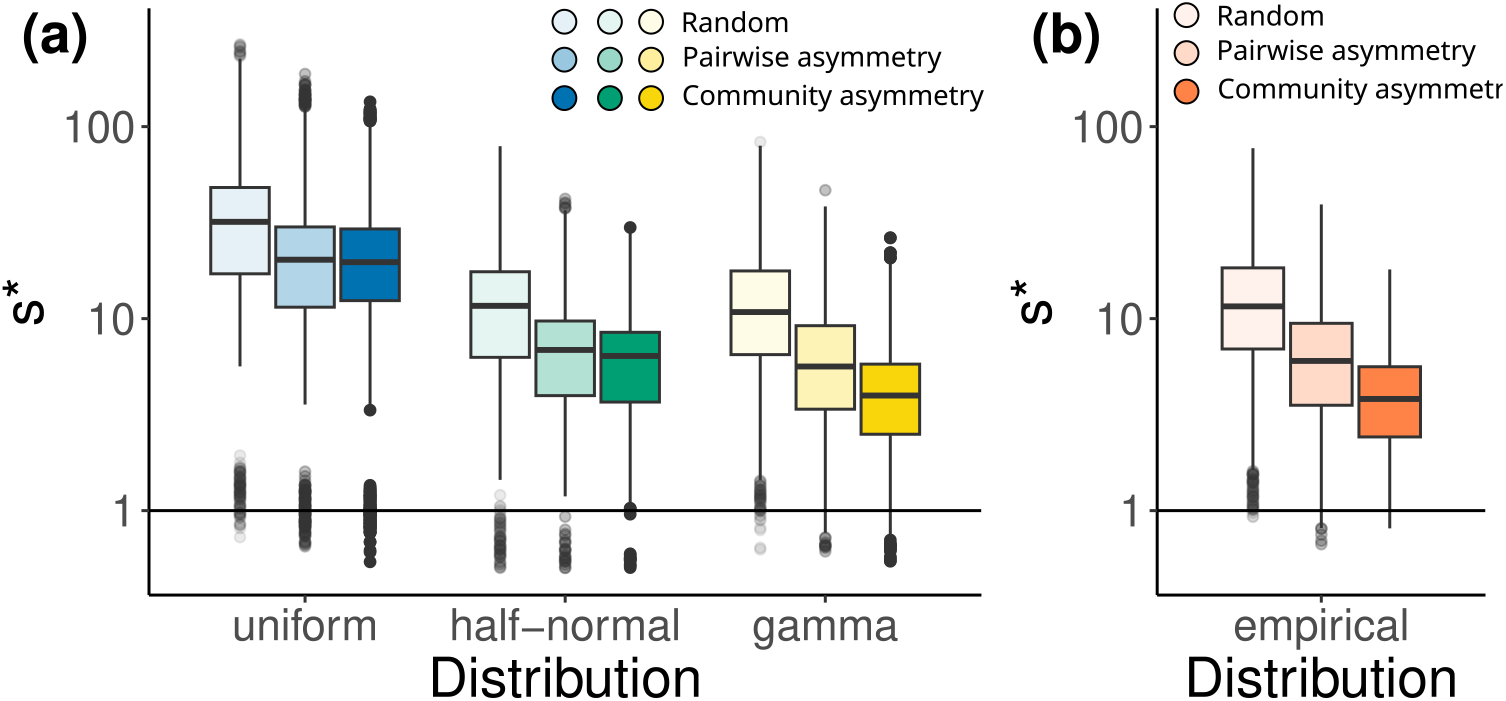
The stabilising effect of asymmetries in theoretical matrices with varying distributions of interaction strengths. We quantify this effect by comparing the stability of randomly patterned and asymmetric matrices for theoretical distributions (a). For comparison, we also quantify the effect in theoretical matrices with the observed, empirical distribution of interaction strengths (b). The level of (in)stability was compared using the critical amount of self-regulation, *s*^*^. As most matrices were unstable, this described how much self-regulation would have to be added to reach stability. Size, connectance and topology of each theoretical matrix was chosen based on one of the 30 empirical systems. They contained interaction strengths drawn from a uniform, normal, or gamma distribution fitted to the same empirical counterparts’ interaction strengths (a) or the observed interaction strengths (b). This was repeated 100 times per empirical data set, resulting in 30 × 100 = 3000 data points per boxplot.

In theoretical matrices with empirical distributions, which we use as a reference, pairwise asymmetric patterns reduced instability and community asymmetry reduced instability even further (Fig. 4 b, in line with the findings of Koch et al., 2023). Correspondingly, for all theoretical distributions, a pairwise asymmetric pattern resulted in a reduction of instability as well. However, adding community asymmetry only had an additional stabilising effect in systems with a gamma distribution. In matrices with half-normal or uniform distributions, there was no additional effect of community asymmetry (Fig. 4 a). This implies that a certain level of skewness is required to enable an additional stabilising effect of community asymmetry. This result was independent of the specific architectures or sizes of the networks, and also held for random topologies (Supplementary Fig. 1) and for larger networks involving hundreds of species (Supplementary Fig. 2).

### 4.3 Skewness enables the stabilising effect of community asymmetry

In a final step, we wanted to obtain a better understanding of how skewness affected the stabilising effect of pairwise and community asymmetry. We created additional sets of matrices using gamma distributions in which we systematically varied the level of skewness, while keeping matrix size *S*, connectance *C* and the mean interaction strength constant. For each level of skewness, we quantified the stabilising effect of asymmetry by calculating stability gain, the ratio of mean *s*^*^ of randomly patterned to asymmetric matrices (Fig. 5 a). We found that for low skewness values between *Ŝ* = 0 and *Ŝ* = 0, stability gain due to pairwise asymmetry and stability gain due to community asymmetry increased equally. An additional stabilising effect of community asymmetry only appeared for higher levels of skewness. For skewness values above *Ŝ* = 2, the stability gain due to pairwise asymmetry remained constant at about 2, while the effect of community asymmetry kept increasing strongly.

**Figure 5:**
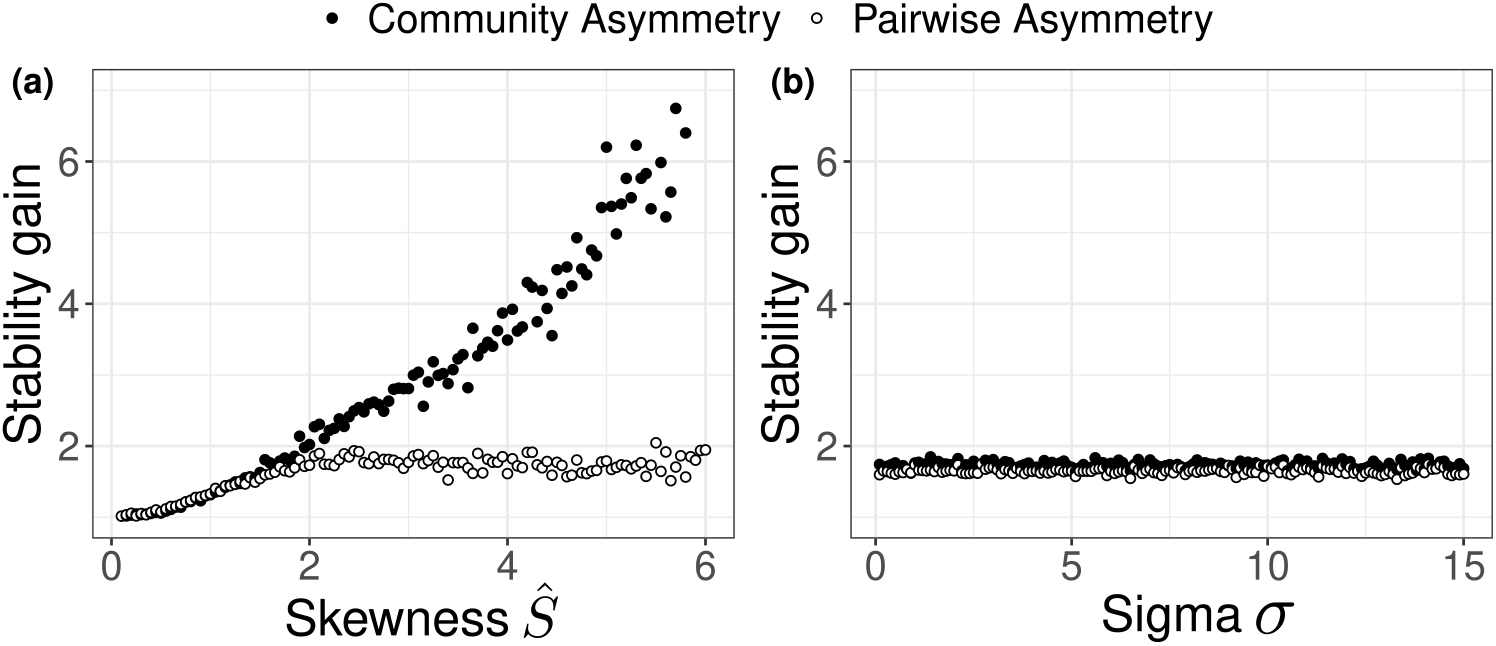
Relation between the shape of interaction strength distribution and the effect of patterning on system stability. Comparison between gamma distributions (a) and half-normal distributions (b). Levels of skewness (in a) and variance (in b) were related to the gain in stability obtained by introducing pairwise and community asymmetric patterning. We calculated stability gain as the mean *s*^*^ of 100 randomly arranged matrices divided by the mean *s*^*^ of 100 asymmetric matrices. Matrix size and connectance were kept constant at *S* = 10 and *C* = 0.8 and the location of non-zero links was chosen randomly. (a) Stability gain in matrices constructed with gamma distributions of varying skewness values. To isolate the effect of skewness, we kept the mean interaction strength constant at −3. To vary the skewness *Ŝ* of a gamma distribution while keeping the mean *m* constant, we calculated the shape *α* and rate *λ* paramters as: *α* = 4*/*(*Ŝ*^2^) and *λ* = *α/m* (b) Stability gain in matrices constructed with half-normal distributions with increasing scale parameters *σ* and thus increasing variance.

In gamma distributions, increasing skewness also increases variance, so that these two properties cannot be seperated. We therefore conducted a similar experiment with half-normal distributions, where variance can be increased but skewness is constant (Fig. 5 b). In this case, we found no relationship between stability gain and the level of variance. Thus, we can conclude that the stabilising effect of community asymmetry indeed depended on skewness, which describes the specific proportion of strong and weak links, and not on variance alone.

## 5 Discussion

Our results show the importance of having many weak and few strong links for enabling stabilising patterns of interaction strengths in competition networks. First, we showed that link strengths derived from empirical competition assemblages have very skewed distributions. Then, we demonstrated that this skewness is necessary to reproduce a stabilising effect of community asymmetry, and hence of hierarchy, which has been found in empirical competition networks (Koch et al., 2023). We did this using theoretical matrices in which asymmetric patterns were generated using random interaction strengths drawn from distributions with varying shapes. We compared uniform, half-normal and gamma distributions and found that a stabilising effect of community asymmetry could only be observed with skewed gamma distributions. By systematically varying the skewness level of gamma distributions, we could confirm that the stabilising effect of community asymmetry indeed depended on skewness.

Competition for space or other resources is often fundamental to realised richness and persistence of biodiversity (Paine, 1966). Yet, our understanding of patterns, mechanisms, drivers and stability underpinning this at assemblages or community level still has significant gaps. One of these gaps is that studies of empirical distributions of competitive interaction strengths are scarce. While there are many studies that quantify the intensity of competition in some way, they tend to focus on individual effects that cannot be easily transferred to the population level link strengths used in ecological network models (Goldberg et al., 1999; Wootton and Emmerson, 2005; Hart et al., 2018). Studies on competitive networks, often in the context of intransitive competition, have therefore been using binary who-beats-whom networks (Laird and Schamp, 2006; Allesina and Levine, 2011; Gallien et al., 2018).

To the best of our knowledge, the only other empirically quantified competitive community matrix, apart from the one published by Koch et al. (2023), is the one presented by Roxburgh and Wilson (2000). In contrast to our results, they identified an approximately uniform distribution of link strength in a single matrix based on a (terrestrial) lawn community. However, their parametrisation was based on experiments with isolated pairs of species rather than on multi-species systems where biomass distributions have naturally formed. In the 30 empirical networks we used for the present study, which were based on observations of whole assemblages (Koch et al., 2023), the skewness we saw in the interaction strengths was largely the result of assemblage dynamics, which generated very skewed distribution of observed species abundances and biomass loss rates (see Supplementary Note 2).

In trophic networks, where there is a long tradition of determining interaction strengths from observations (Paine, 1992; Polis, 1994; Ruiter et al., 1995; Wootton, 1997; Berlow, 1999; Emmerson and Yearsley, 2004; Neutel, Heesterbeek, Koppel, et al., 2007; Neutel and Thorne, 2014), there is ample evidence of skewed interaction strengths across various types of environments (Ruiter et al., 1995; Berlow et al., 2004; Wootton and Emmerson, 2005; Jacquet et al., 2016). The “few strong, many weak links” distribution has also been observed in mutualistic systems (Jordano, 1987; Bascompte, Jordano, et al., 2006). Even though the exact methods and definitions for quantifying interaction strengths often differ, skewed interaction strengths are a consistent observation and thus appear to be a common property of natural ecological networks.

Our result that the effect of hierarchy on stability only appeared with skewed distributions of interaction strengths was found to hold for random topologies and for larger theoretical matrices with several hundred species (Supplementary Figs. 1 and 2). This is interesting, as previous results based on very large, unstructured matrices indicate that the exact shape of the distribution of interaction strengths does not influence stability (Allesina and Tang, 2012; Allesina and Tang, 2015). For these matrices, only the mean and the variance of matrix elements affect stability, while higher moments of the distribution, like the skewness, are not relevant (Tao et al., 2010). Our findings indicate that the proportion of strong and weak links can have an important effect on stability when we look at matrices with non-random patterns.

Our results can be understood by looking at the feedback structure of the systems. In general, pairwise asymmetry is stabilising, as it reduces the amplifying effect of positive 2-link loops. In contrast to this, community asymmetry is stabilising as it reduces the strength of long, negative feedback loops (Koch et al., 2023). If these long, negative feedback loops are too strong compared to shorter loops, the excessive negative feedback can cause oscillatory instability (Levins, 1974). Community asymmetry avoids this destabilising imbalance, which explains its additional stabilising effect compared to pairwise asymmetry. As we found that the stabilising effect of community asymmetry only appeared when interaction strengths were skewed, our results indicate that this imbalance can only form when there is a sufficient level of skewness. Further analysis of feedback loop strengths would be necessary to confirm this and to uncover the exact conditions that lead to the emergence of oscillatory instability.

The contrasting mechanisms that explain the stabilising effects of pairwise and community asymmetry mean that we have to consider two “regimes” of instability, one governed by positive feedback, the other governed by excessive negative feedback (Levins, 1974). These two regimes are not only relevant to competitive systems, but may also play a role in food webs. In fact, the idea that instability is caused by overshoots leading to unstable oscillations is common in food web literature (McCann et al., 1998; Emmerson and Yearsley, 2004; Gellner and McCann, 2016), although it is usually not linked to the concept of feedback loops.

In conclusion, we showed that similar to what has been found in food webs and mutualistic systems, competition networks contain many weak and few strong links. We furthermore demonstrated that this type of distribution enables stabilising patterns in competition webs. These insights have important implications for theoreticians exploring structure in random matrices, which often use normal or uniform distributions of matrix elements (Emmerson and Yearsley, 2004; Allesina and Tang, 2012). Our study shows that when specific patterns of interaction strengths are introduced, the shape of distributions can have an effect on stability. This needs to be considered, when we want to explore and understand stabilising mechanisms in real systems, which typically have very skewed interaction strengths.

## 6 Declarations

## Acknowledgements

We thank the Plant Ecology Group at the University of Tübingen, led by Katja Tielbörger, for office space, moral support and helpful discussions. Specifically, we would like to thank Pierre Liancourt for his suggestion to analyse the skewness-kurtosis relationship of interaction strength distributions. Additionally, we thank Jon Pitchford and the Complexity-Stability Reading Group at the University of York for insightful discussions of the results.

## Author contributions

F.K. conceived the study and carried out the initial analysis. DKAB provided empirical datasets. FK, AMN and KTA designed further analysis steps and discussed the results. FK wrote the first manuscript draft, with input from KTA and AMN. All authors contributed critically to revisions.

