## Supplementary Material for "Many weak and few strong links: The importance of interaction strength distributions for stabilising patterns in competition networks"

### 1 Supplementary Note 1: Derivation of interaction strengths

The interaction strengths were derived from an observational data set of overgrowth competition between bryozoan colonies (Koch et al., 2023). For clarity and background, we describe below the methods described in Koch et al., 2023. The data sets originate from 3 different polar regions: Signy Island and Adelaide Island in the Antarctic as well as Spitzbergen, Svalbard in the Arctic. For each sampling location, the data set includes an abundance table, containing the number of colonies per species, as well as a species-contact-matrix that summarises the outcomes of all spatial contests between each pair of competing species. For details on data collection and preparation, see Barnes and Kuklinski, 2004 and Koch et al., 2023, respectively.

The observed competitive outcomes were first translated into energy loss rates $f_{ij}$, describing the loss in biomass of a species $i$ as a result of the interference competition with species $j$. Total energy loss rate [biomass/time] was calculated by summing up the number of wins $W_{ij}$, losses $L_{ij}$ and draws $D_{ij}$, each weighted with a cost value, which describe a fixed proportion of biomass loss per colony per time:

$$f_{ij} = -0.1W_{ij} - 0.9L_{ij} - 0.2D_{ij} \tag{1}$$

Then, the energy loss rates were used to calculate interaction strengths, which are defined as the elements of the Jacobian matrices. These elements $a_{ij}$ described the per-capita effect of a change in the biomass of species $j$ on the biomass of species $i$. This step was based on a steady-state assumption, which was made to be able to test network stability. Furthermore, it was assumed that the dynamics around this steady state were described by the following Lotka-Volterra equations:

$$\frac{dX_i}{dt} = r_i X_i - \sum_{j=1}^{n} c_{ij} X_i X_j \tag{2}$$

where $X_i$ was the population density of species $i$, $r_i$ its intrinsic growth rate, and $c_{ij}$ a constant which described competition intensity between species $i$ and $j$. The interaction strengths were the partial derivatives of these equations, evaluated at equilibrium. For interspecific interactions, the interaction strength is thus $a_{ij} = -c_{ij}X_i^*$. For intraspecific interactions, they are $a_{ii} = r_i - \sum_{ij}^{n} c_{ij}X_j^* - 2c_{ii}X_i^*$, which simplifies into $a_{ii} = -c_{ii}X_i^*$ at equilibrium where $r_i - \sum_{ij}^{n} c_{ij}X_j^* - c_{ii}X_i^* = 0$.

Energy loss rates $f_{ij}$ were assumed to correspond to the loss term due to competition $-c_{ij}X_iX_j$ and the observed abundance $B_i$ of species $i$ was assumed to correspond to its equilibrium density. Therefore, the interaction strengths could be calculated as:

$$a_{ij} = -c_{ij}X_i^* = \frac{-c_{ij}X_i^*X_j^*}{X_j^*} = \frac{f_{ij}}{B_j} \tag{3}$$

In a final step the interaction strengths were normalised, following Neutel and Thorne, 2014, by dividing each matrix element by the absolute values of its corresponding diagonal term: $\bar{a_{ij}} = \frac{a_{ij}}{|a_{ii}|}$. Missing diagonal elements were estimated as the mean interaction strengths, multiplied with $-0.1$(Koch et al., 2023). The stability properties of matrices with varying diagonal elements are difficult to compare. The normalisation makes such comparisons possible, as it translates the matrices' diagonal structure into its off-diagonal structure and results in all diagonal terms being -1, just as in May-type random matrices (see also Thorne et al., 2021).

#### 2 Supplementary Note 2: Skewness in biomass distribution

The skewed distribution of interaction strengths we found in empirical data can be seen as a consequence of a colonisation-competition trade-off in bryozoan assemblages (Barnes and Conlan, 2007). Assemblages contain both strong colonisers who tend to dominate in biomass (Barnes and Clarke, 1998) but lose most contests, and strong competitors who are less abundant but win most contests.

Table 1 illustrates these strong differences in abundance within one data set. As shown in Figure 1 A for the same example system, this resulted in very skewed energy loss rates. Translating energy loss rates to interaction strengths then further increased the level of skewness because energy loss rates were divided by abundances, which are also skewed (Figure 1 B). Finally, the normalisation of interaction strengths slightly reduced the level of skewness again (Figure 1 C). Fig 1 D shows the skewness-kurtosis relationship of energy loss rates, raw and normalised interaction strengths in all data sets, demonstrating that what is shown in the example in Figure 1 A-C applies to all 30 networks. The skewness that we saw in the interaction strengths was thus an inherent property of the empirical data and was not imposed by the derivation of interaction strengths.

Table 1: Abundance [number of colonies] for each species in example data set Rothera 3

| Species | Abundance |
|---|---:|
| H | 3 |
| M | 5 |
| S | 7 |
| L | 8 |
| E | 9 |
| C | 9 |
| Ca | 12 |
| Ar | 17 |
| X | 20 |
| Ai | 39 |
| F | 820 |

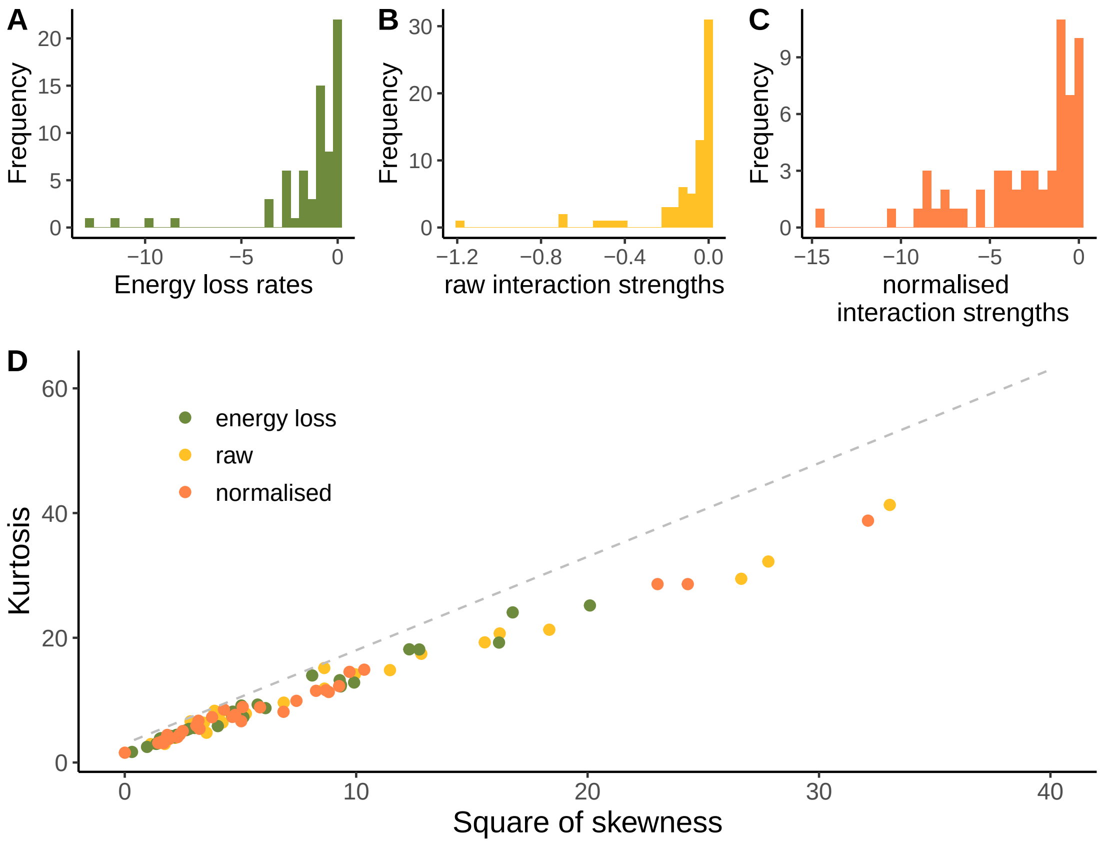


Figure 1: **Distribution of energy loss rates, raw and normalised interaction strengths**. (A-C) histograms showing the distribution of energy loss rates, raw interaction strengths and normalised interaction strengths for example data set *Rothera 3* (D) Skewness-kurtosis relationship of energy loss rates, raw and normalised interaction strengths for all data sets.

### 3 Supplementary Tables

Supplementary Table 1: **Properties of empirical interaction strengths** For each data set we show the number of species, $S$, and the connectance $C$, which is the number of realised links relative to the number of possible links. Furthermore, distribution shape is characterised by their skewness $\hat{S}$, describing how asymmetric the distribution of interaction strengths is, and kurtosis $\hat{K}$ which describes how peaked the distribution is.

| Name | Number of species $S$ | Connectance $C$ | Skewness $\hat{S}$ | Kurtosis $\hat{K}$ |
|---|---|---|---|---|
| Rothera_1 | 11 | 0.45 | -4.80 | 28.61 |
| Rothera_2 | 9 | 0.47 | -1.51 | 4.08 |
| Rothera_3 | 11 | 0.55 | -1.36 | 4.42 |
| Signy_1 | 8 | 0.89 | -1.54 | 4.49 |
| Signy_2 | 8 | 0.89 | -1.78 | 6.70 |
| Signy_3 | 8 | 0.89 | -2.42 | 8.85 |
| Signy_4 | 8 | 0.93 | -3.22 | 14.88 |
| Spitzbergen_H1_12_1 | 8 | 0.93 | -3.12 | 14.53 |
| Spitzbergen_H1_12_2 | 8 | 0.96 | -2.16 | 7.31 |
| Spitzbergen_H1_6_1 | 7 | 0.95 | -2.62 | 8.13 |
| Spitzbergen_H1_6_2 | 7 | 0.95 | -4.93 | 28.60 |
| Spitzbergen_H2_12_1 | 5 | 0.80 | -0.06 | 1.57 |
| Spitzbergen_H2_12_2 | 6 | 0.73 | -2.72 | 9.89 |
| Spitzbergen_H2_6_1 | 5 | 0.60 | -1.30 | 3.12 |
| Spitzbergen_H2_6_2 | 6 | 0.73 | -2.95 | 11.58 |
| Spitzbergen_H3_12_1 | 6 | 0.93 | -1.20 | 3.14 |
| Spitzbergen_H3_12_2 | 7 | 0.86 | -1.58 | 5.04 |
| Spitzbergen_H3_6_1 | 7 | 1.00 | -1.76 | 5.91 |
| Spitzbergen_H3_6_2 | 7 | 0.95 | -2.18 | 7.63 |
| Spitzbergen_K1_12_1 | 9 | 0.83 | -5.67 | 38.78 |
| Spitzbergen_K1_12_2 | 6 | 0.93 | -1.94 | 7.26 |
| Spitzbergen_K1_6_1 | 10 | 0.93 | -2.07 | 8.49 |
| Spitzbergen_K1_6_2 | 10 | 0.93 | -2.26 | 8.90 |
| Spitzbergen_K2_12_1 | 6 | 0.87 | -3.04 | 12.25 |
| Spitzbergen_K2_12_2 | 4 | 0.83 | -2.24 | 6.62 |
| Spitzbergen_K2_6_1 | 7 | 0.86 | -2.87 | 11.50 |
| Spitzbergen_K2_6_2 | 7 | 0.86 | -1.39 | 3.79 |
| Spitzbergen_K3_12_1 | 5 | 0.90 | -2.97 | 11.31 |
| Spitzbergen_K3_12_2 | 5 | 0.90 | -1.27 | 3.37 |
| Spitzbergen_K3_6_2 | 6 | 0.93 | -1.80 | 5.41 |

Supplementary Table 2: **Fitted parameter values for theoretical distributions** Theoretical distributions were fitted to each empirical set of interaction strengths using the R package *fitdistrplus*.

| Name | uniform | | half-normal | gamma | |
|---|---|---|---|---|---|
| | min | max | scale $\sigma$ | shape $\alpha$ | rate $\lambda$ |
| Rothera_1 | -76.00 | -0.0037 | 11.51 | 0.39 | 0.08 |
| Rothera_2 | -18.01 | -0.0095 | 4.80 | 0.44 | 0.12 |
| Rothera_3 | -14.58 | -0.0296 | 4.40 | 0.72 | 0.24 |
| Signy_1 | -9.24 | -0.0402 | 2.85 | 0.74 | 0.36 |
| Signy_2 | -10.18 | -0.0271 | 2.58 | 0.75 | 0.40 |
| Signy_3 | -13.19 | -0.0545 | 3.21 | 0.78 | 0.36 |
| Signy_4 | -24.94 | -0.0373 | 4.66 | 0.58 | 0.23 |
| Spitzbergen_H1_12_1 | -40.74 | -0.0122 | 7.61 | 0.42 | 0.11 |
| Spitzbergen_H1_12_2 | -32.95 | -0.0318 | 8.53 | 0.55 | 0.11 |
| Spitzbergen_H1_6_1 | -32.68 | -0.0038 | 7.81 | 0.35 | 0.09 |
| Spitzbergen_H1_6_2 | -48.09 | -0.0137 | 6.80 | 0.39 | 0.13 |
| Spitzbergen_H2_12_1 | -1.42 | -0.0829 | 0.45 | 2.08 | 2.74 |
| Spitzbergen_H2_12_2 | -16.81 | -0.1950 | 2.77 | 0.85 | 0.33 |
| Spitzbergen_H2_6_1 | -11.38 | -0.0298 | 2.25 | 0.52 | 0.17 |
| Spitzbergen_H2_6_2 | -18.49 | -0.0944 | 2.86 | 0.63 | 0.26 |
| Spitzbergen_H3_12_1 | -9.40 | -0.0376 | 2.70 | 0.92 | 0.33 |
| Spitzbergen_H3_12_2 | -15.60 | -0.0141 | 4.08 | 0.88 | 0.24 |
| Spitzbergen_H3_6_1 | -26.31 | -0.0191 | 7.02 | 0.66 | 0.12 |
| Spitzbergen_H3_6_2 | -50.06 | -0.0295 | 10.92 | 0.63 | 0.08 |
| Spitzbergen_K1_12_1 | -154.02 | -0.0520 | 22.56 | 0.42 | 0.05 |
| Spitzbergen_K1_12_2 | -49.16 | -0.0003 | 9.44 | 0.39 | 0.05 |
| Spitzbergen_K1_6_1 | -32.79 | -0.0231 | 8.80 | 0.64 | 0.14 |
| Spitzbergen_K1_6_2 | -38.18 | -0.0243 | 10.39 | 0.62 | 0.12 |
| Spitzbergen_K2_12_1 | -86.04 | -0.0079 | 13.57 | 0.31 | 0.03 |
| Spitzbergen_K2_12_2 | -39.78 | -0.0222 | 5.47 | 0.35 | 0.05 |
| Spitzbergen_K2_6_1 | -49.94 | -0.0007 | 8.89 | 0.34 | 0.07 |
| Spitzbergen_K2_6_2 | -28.96 | -0.0002 | 7.68 | 0.41 | 0.07 |
| Spitzbergen_K3_12_1 | -68.06 | -0.0016 | 9.85 | 0.31 | 0.04 |
| Spitzbergen_K3_12_2 | -36.22 | -0.0010 | 7.39 | 0.34 | 0.04 |
| Spitzbergen_K3_6_2 | -47.94 | -0.0041 | 10.10 | 0.39 | 0.05 |

### 4 Supplementary Figures

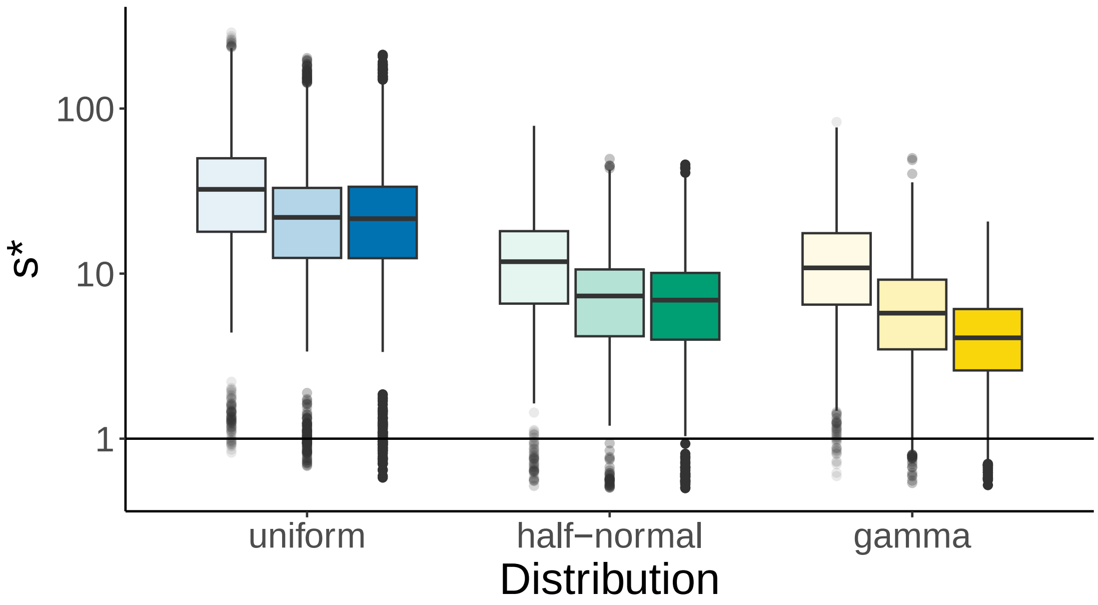


Supplementary Figure 1: **Stabilising effect of asymmetries in matrices with varying distributions of interaction strengths and random topology.** Comparison of stability in random (lightest shade), pairwise asymmetric (medium shade) and community asymmetric (darkest shade) matrices with varying distributions of interaction strengths. Stability is measured as the critical amount of self-regulation $s^*$. Matrices are created exactly as those shown in Fig 3 A of the main analysis, but instead of using empirical topologies, the location of non-zero links is chosen at random.

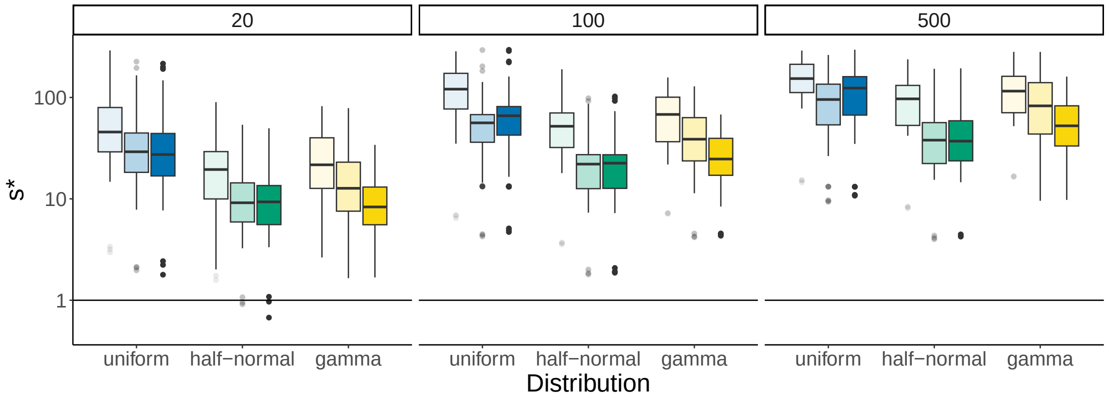


Supplementary Figure 2: **Stabilising effect of asymmetries in larger matrices**. Comparison of stability in random (lightest shade), pairwise asymmetric (medium shade) and community asymmetric (darkest shade) matrices with varying distributions of interaction strengths and varying sizes. Matrices were created using the same distributions as in the main analysis. Connectance was chosen corresponding to the empirical matrices, but size was varied. Increasing the size of the matrices led to an overall increase in instability. In matrices with gamma- and normally distributed interaction strengths, the stability effects of patterning remained the same for all sizes: For gamma, community asymmetry was more stable than pairwise asymmetry, while in the normal distribution, we see no difference between the two. In the large uniformly distributed matrices, we observed that community asymmetry increases instability compared to pairwise asymmetry.
